# Signatures of Selection in the Human Antibody Repertoire: Selective Sweeps, Competing Subclones, and Neutral Drift

**DOI:** 10.1101/145052

**Authors:** Felix Horns, Christopher Vollmers, Cornelia L. Dekker, Stephen R. Quake

## Abstract

Antibodies are created and refined by somatic evolution in B cell populations, which endows the human immune system with the ability to recognize and eliminate diverse pathogens. However, the evolutionary processes that sculpt antibody repertoires remain poorly understood. Here, using an unbiased repertoire-scale approach, we show that the molecular signatures of evolution are evident in human B cell lineages and reveal how antibodies evolve somatically. We measured the dynamics and genetic diversity of B cell responses of five adults longitudinally before and after influenza vaccination using high-throughput antibody repertoire sequencing. We identified vaccine-responsive B cell lineages that carry signatures of selective sweeps driven by positive selection, and discovered that they often display evidence for selective sweeps favoring multiple subclones. We also found persistent B cell lineages that exhibit stable population dynamics and carry signatures of neutral drift. By exploiting the linkage between B cell fitness and antibody binding affinity, we demonstrated the potential for using signatures of selection to identify antibodies with high binding affinity. This quantitative characterization reveals that antibody repertoires are shaped by an unexpectedly broad spectrum of evolutionary processes and shows how signatures of evolutionary history can be harnessed for antibody discovery and engineering.

**One Sentence Summary:** Molecular signatures of somatic evolution reveal that diverse evolutionary processes ranging from strong positive selection to neutral drift sculpt human antibodies.

## Main Text

Antibodies are created through evolutionary processes involving mutation and selection, all of which unfold in B cell populations. As proposed by Burnet in his “clonal selection theory” in 1957, the concepts of population genetics offer an avenue for understanding how antibody repertoires evolve (*1*). Yet, after 60 years of progress in immunology, the somatic evolution of human antibodies remains poorly understood and immunology has yet to benefit from the quantitative theories and models of population genetics which have been transformative in our understanding of evolution at the organismal level.

Selective processes are widely thought to exist during affinity maturation, as they would help focus the antibody repertoire on antibodies that can bind antigens with high affinity (*2*–*4*). After infection or immunization, activated B cells migrate to germinal centers (GCs) where they undergo genetic diversification via somatic hypermutation and selection for affinity-enhancing mutations. Within several weeks after antigenic challenge, this Darwinian process generates antibodies with increased average affinity to the antigen (*5*, *6*). Despite intense experimental effort focused on the cellular and molecular mechanisms of affinity maturation (*7*–*12*), the evolutionary process itself remains poorly characterized. Each GC is founded by tens to hundreds of distinct B cell clones and this diversity is often lost due to competition between clones as affinity maturation proceeds (*13*). However, how the competition unfolds between the genetically diverse variants within the same clonal B cell lineage has not been described, despite its importance for the emergence of protective antibodies. Furthermore, although it is often presumed that the same evolutionary processes affect B cells across the entire repertoire, some B cell types, such as B-1 cells, do not participate in classical affinity maturation, and little is known about the full diversity of evolutionary patterns that shape human antibody repertoires.

Here, we characterize the dynamics and somatic evolution of human B cell lineages using high-throughput sequencing of the antibody repertoire and analytical methods inspired by population genetics. We performed time-resolved measurements of antibody repertoires in healthy young adults before and after seasonal influenza vaccination. We identified vaccine-responsive B cell lineages that expanded dramatically after vaccination, and we show that patterns of genetic variation within these lineages reflect a history of strong positive selection. This selection drove recurrent selective sweeps during somatic evolution, and many vaccine-responsive B cell lineages display evidence for selective sweeps favoring multiple subclones. Other abundant B cell lineages are stable and lack a response to vaccination; we show that these lineages carry signatures of neutral evolution. Finally, we present an approach for using phylogenetic information to rapidly identify potential high-affinity antibodies and affinity-enhancing mutations. Our results offer a detailed portrait of the somatic evolutionary processes that shape human antibody repertoires and link models of evolution with quantitative measurements of the human immune system.

We measured the dynamics of the antibody repertoires of five healthy young adults before and after vaccination in late spring 2012 with the 2011–2012 trivalent seasonal flu vaccine (Figure 1A). Volunteers were influenza vaccine-naïve for the 2010–2011 and 2011–2012 influenza seasons. We sampled peripheral blood at the time of vaccination and 1, 4, 7, 9, and 11 days afterwards (D0, D1, D4, D7, D9, and D11), as well as 3 and 5 days before vaccination (D-3 and D-5). We sequenced transcripts of the immunoglobulin heavy chain gene (IGH) using RNA extracted from peripheral blood mononuclear cells (Materials and Methods). The sequences span ~100 bp of the variable region including complementarity determining region 3 (CDR3), enabling tracking of the dynamics of clonal B cell lineages. We used molecular barcoding to mitigate errors arising during library preparation and sequencing, enabling accurate measurement of genetic diversity (*14*).

**Figure 1.**
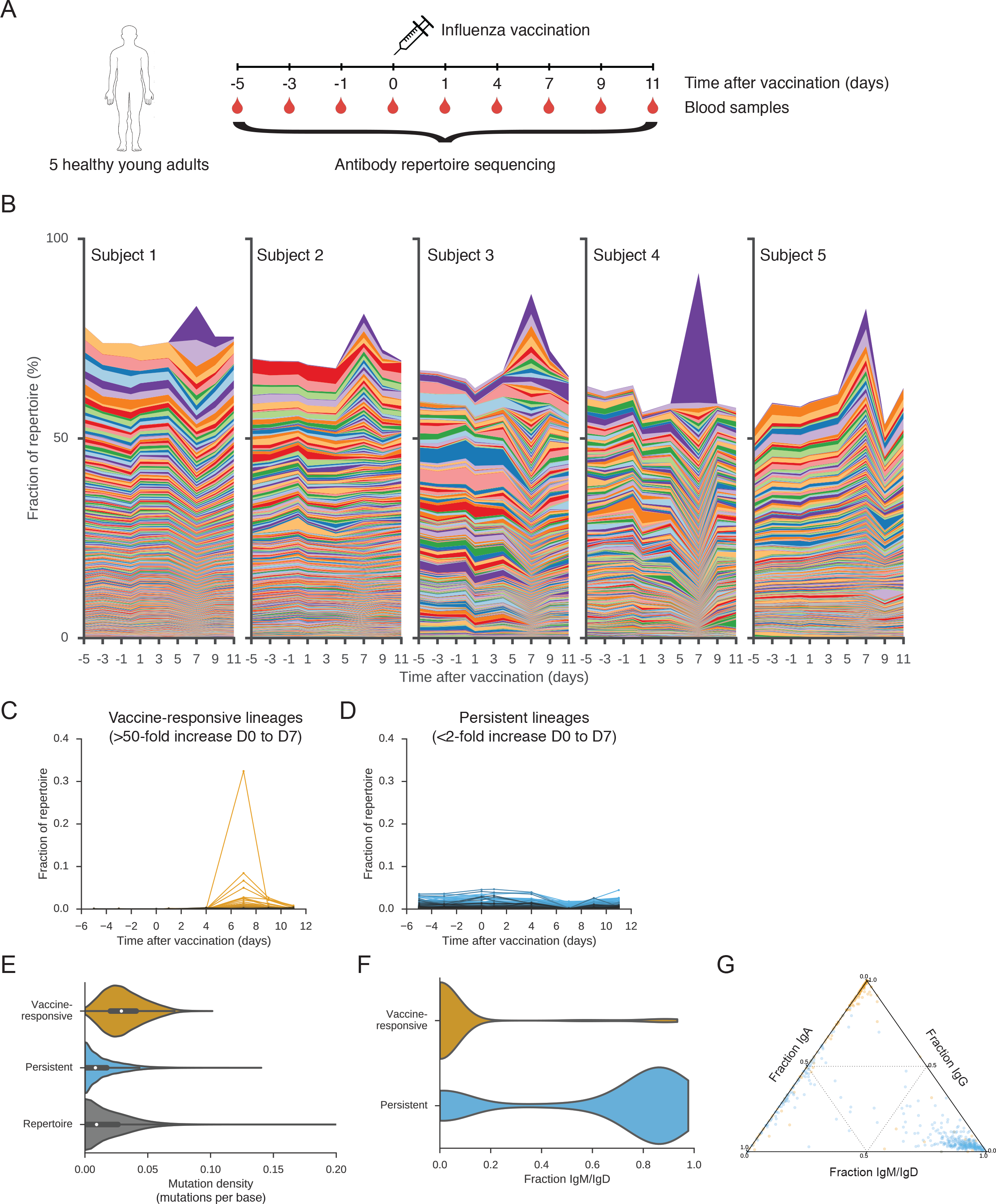
Dynamics and molecular features of antibody repertoires. (A) Schematic of experiment design. (B) Dynamics of antibody repertoires. Each line represents a clonal B cell lineage and its width indicates the fractional abundance of that lineage (the number of unique sequences belonging to the lineage divided by the number of unique sequences in the entire repertoire) at a given time. Lineages that compose >0.01% of the repertoire at D7 are shown. (C) Dynamics of vaccine-responsive lineages. (D) Dynamics of persistent lineages. In (C) and (D), each line represents a clonal lineage. (E) Distributions of somatic mutation density within the V gene in sequences belonging to vaccine-responsive lineages, persistent lineages, or the entire antibody repertoire. Mutations were called by comparison with the germline sequence. (F) Distributions of the fraction of sequences within each clonal lineage that were the IgM or IgD isotypes among vaccine-responsive and persistent lineages. (G) Fractions of sequences in each clonal lineage that were IgM or IgD, IgG, or IgA. Each dot is a lineage and is positioned according to the isotype composition of that lineage and colored according to identification as vaccine-reponsive (yellow) or persistent (blue).

To identify sequences that belong to the same clonal lineage, defined as those that share a common naïve B cell ancestor, we first grouped sequences having the same V and J germline genes and CDR3 length. Within each group, we identified clonal lineages by performing single-linkage clustering on the CDR3 sequence using a cutoff of 90% sequence identity, an approach that accurately partitions sequences into clones (*15*, *16*).

To visualize how the composition of the antibody repertoire changed after vaccination, we examined the fractional abundance of clonal B cell lineages over time (Figure 1B). All five subjects had a strong response to vaccination, exhibiting dramatic changes in the relative abundance of B cell lineages within 7 days, which is characteristic of a memory recall response to vaccination (*14*). In each subject’s repertoire, we identified 36 ± 12 (mean ± s.d., range 16 – 49) B cell lineages that expanded >50-fold between D0 and D7 after vaccination (Figure 1C; Table S1). In contrast, across a similar timespan in the absence of vaccination (between D0 and D-5), only 6 ± 4 lineages within each subject expanded to this extent (Figure S1A) and expansion of these lineages may be attributable to exposure to environmental antigens. Because most of these “vaccine-responsive” lineages were undetectable prior to vaccination, the identification of vaccine-responsive lineages was robust to the specific choice of fold-change cutoff (Figure S1A). Together, these vaccine-responsive lineages accounted for 22% ± 12% (mean ± s.d., range 10% – 43%) of each subject’s repertoire during peak response at D7. Vaccine-responsive antibodies have high levels of somatic mutation (Figure 1E) and are predominantly class-switched (Figure 1F and Figure 1G), as expected for memory B cells. Thus, influenza vaccination triggers rapid memory recall of dozens of clonal B cell lineages in healthy human adults.

We discovered that each subject harbored a distinct set of clonal B cell lineages that exhibited high abundance throughout the study and were unresponsive to vaccination (Figure 1D). In each subject, we detected 83 ± 23 (mean ± s.d., range 44 – 111) of these “persistent lineages”, which together accounted for 22% ± 8% (mean ± s.d., range 10% - 33%) of the repertoire at any time point (Table S1). Persistent lineages displayed remarkably stable dynamics compared with vaccine-responsive lineages (Figure S1B), implying detailed balance in their cellular population dynamics and mRNA expression levels. Persistent antibodies have low levels of somatic mutation (Figure 1E) and are mostly the IgM isotype, but a minority of persistent lineages is composed predominantly of the IgA isotype (Figure 1F and Figure 1G). Thus, many human antibody repertoires possess a large complement of persistent B cell lineages, which have slow turnover and do not respond dynamically to influenza vaccination.

Evolutionary history leaves enduring signatures in the genetic diversity of populations. Vaccine-responsive B cell lineages carrying memory B cells underwent affinity maturation when the subjects were exposed to influenza antigens for the first time. We reasoned that examination of the patterns of genetic variation within these lineages might give insight into the evolutionary processes that unfolded during affinity maturation. Visualizing the phylogenies of clonal B cell lineages revealed that many vaccine-responsive lineages possess highly imbalanced branching structure across many levels of depth, suggesting that these lineages experienced recurrent selective sweeps (Figure 2A). This signature reflects continuous adaptive evolution under strong positive selection and has been found in many asexual populations evolving under sustained adaptive pressure, such as influenza virus (*17*) and human immunodeficiency virus (*18*).

**Figure 2.**
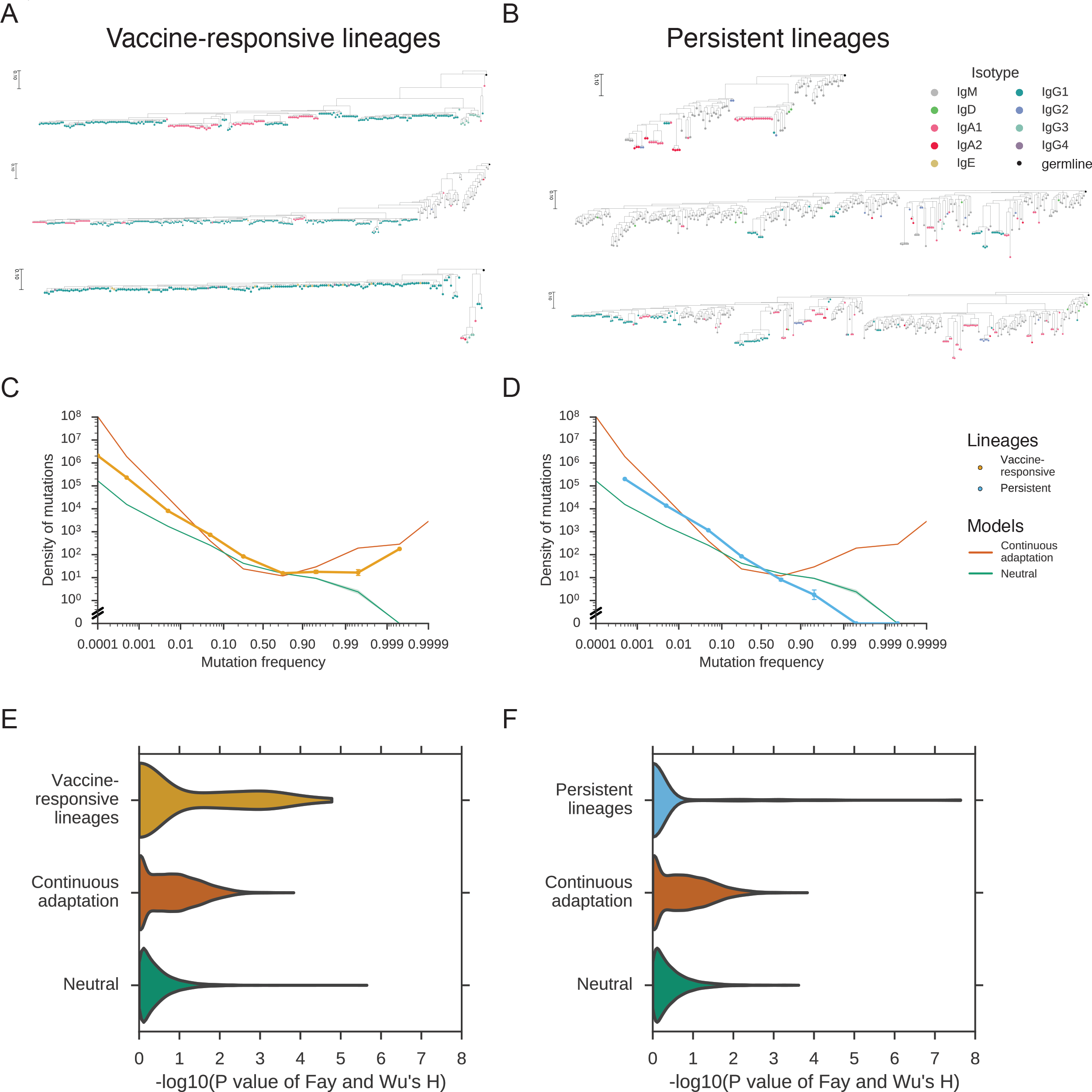
Genetic signatures of somatic evolution in clonal antibody lineages. (A and B) Examples of phylogenies of vaccine-responsive (A) or persistent (B) clonal B cell lineages. Leaves are colored by isotype. Phylogenies are rooted on the germline sequence. (C and D) Site frequency spectrums (SFSs) averaged across all vaccine-responsive lineages (C) or persistent lineages (D). Error bars indicate standard error of the mean. SFSs generated by population genetic models of continuous adaptation driven by strong positive selection (orange) and neutral drift (green) are shown for comparison. Shading indicates standard error of the mean across simulations (100 replicates). (E and F) Distribution of significance scores of Fay and Wu’s H statistic for vaccine-responsive lineages (E) or persistent lineages (F) compared against models of neutral evolution and continuous adaptation driven by strong selection. Distributions for the models were generated via simulations of evolving populations having sizes sampled from the observed population size distributions of vaccine-responsive or persistent lineages (10,000 replicates).

To quantitatively characterize the evolutionary histories of clonal B cell lineages, we examined the frequency spectrum of derived somatic mutations, also known as the site frequency spectrum (SFS). The SFS carries detailed information about evolutionary
history and can be useful for detecting selective processes. In continuously adapting asexual populations, the SFS exhibits a distinct excess of high frequency variants, which can be used to rule out neutral models and infer positive selection (*19*), as in the cases of influenza virus (*17*) and human immunodeficiency virus (*18*). We calculated the SFS of each clonal B cell lineage based on somatic point mutations relative to the personalized germline V and J gene sequences for each subject because the ancestral state is known with high confidence for these sites (Figure S1C; Materials and Methods). We compared the observed SFSs against population genetic models of neutral evolution [Kingman coalescent (*20*)] and continuous adaptation [Bolthausen-Sznitman coalescent (*21*)] using computer simulations (Materials and Methods).

We first visualized the SFS as an average over all vaccine-responsive lineages and found that the SFS was highly skewed, exhibiting a large excess of high frequency somatic mutations in clear disagreement with the neutral model (Figure 2C). Instead, the model of positive selection has an excellent fit to the data, implying that the dominant mode of evolution in vaccine-responsive lineages is recurrent selective sweeps driven by the occurrence of beneficial mutations. This finding is consistent with the classical model of affinity maturation: affinity-enhancing mutations arise and selection focuses the repertoire on these variants, driving the loss of intraclonal diversity. The presence of deep branches harboring persistent minor alleles within each clonal lineage reveals that memory B cells frequently exit GCs while selection continues, preventing complete loss of diversity due to selective sweeps. These characteristics were also observed in the SFS averaged over vaccine-responsive lineages in each subject (data not shown), indicating that signatures of positive selection within vaccine-responsive lineages are a conserved feature of healthy human immune systems.

Next, we sought to characterize the patterns of somatic evolution at the resolution of individual clonal B cell lineages. While individual lineages have fewer somatic mutations and exhibit sparse spectra compared to the population averages, we found that many vaccine-responsive lineages have a large excess of high frequency mutations (Figure S2A). To quantitatively detect selection, we used Fay and Wu’s H statistic (*22*), which was originally devised to detect high-frequency hitchhiking alleles that are transiently associated with selective sweeps in recombining populations but can also sensitively detect selective sweeps in asexual populations. Using H, we found that 32% of vaccine-responsive lineages deviate significantly from the neutral model (Figure 2E, Figure S2B, and Figure S2C; P < 0.05). We also directly measured the non-monotonicity of the SFS, and found that 14% of vaccine-responsive lineages deviated significantly from neutrality by this alternative metric for selection (Figure S2D and Figure S2E). Nearly every subject had at least one vaccine-responsive lineage that evidently experienced selection (Figure S2G), and the failure to detect selection in every vaccine-responsive lineage is consistent with statistical limits of detection arising from population sizes of the lineages (Figure S2F). Indeed, selection was detected at a rate that is consistent with a model in which every vaccine-responsive lineage evolved under strong positive selection (Figure 2E and Figure S2H), suggesting that affinity maturation forges the entire vaccine-responsive antibody repertoire. Further, high-frequency derived mutations are enriched within complementarity determining regions (CDRs), which form the antibody-antigen binding interface and often evolve under positive selection (*23*, *24*); such mutations are depleted in framework regions (FWRs; Figure S2I), which form the structural scaffold of the antibody molecule and typically evolve under purifying selection (*23*, *24*). Together these observations demonstrate that evolutionary history can be quantitatively characterized at the resolution of individual clonal B cell lineages and support the conclusion that vaccine-responsive lineages evolved under strong positive selection for antibody-antigen interactions.

Persistent antibody lineages have a strikingly different mode of evolution. When we visualized the SFS as an average over all persistent lineages, we found that its shape is consistent with neutral evolution, lacking an excess of high frequency somatic mutations (Figure 2D). Indeed, persistent lineages had no mutations at frequencies above 99%, in agreement with the prediction of the neutral model but not the model of positive selection. This pattern was also clearly evident in individual clonal lineages (Figure S3A), as reflected in their balanced phylogenies, which are characteristic of neutral drift-like evolution (Figure 2B). Using Fay and Wu’s H statistic, we found that nearly every persistent lineage (94%) had no significant departure from neutrality (Figure 2F, Figure S3B, and Figure S3C; P > 0.05). We also found no significant departure from neutrality for nearly every persistent lineage (99%) using the non-monotonicity of the SFS as a metric for selection (Figure S3D and Figure S3E). Persistent lineages had large population sizes comparable to those of vaccine-responsive lineages (100 to ~11,000 sequences; Figure S3F), indicating that limits of detection arising from population size cannot explain the failure to detect selection. Indeed, the rate at which we detected selection on persistent lineages is much lower than the detection limit (Figure S3H). Thus, persistent lineages evolve in a manner consistent with neutrality, indicating that neutral birth-death processes are responsible for the expansion and maintenance of a substantial fraction of the antibody repertoire.

The molecular features of persistent lineages are characteristic of B-1 cells, a B cell subtype that has a different life history than the better-studied B-2 cells. Both persistent lineages and B-1 cells are mostly IgM (*25*) with a minority of lineages composed predominantly of IgA (*26*) (Figure 1F and Figure 1G), and have low levels of somatic hypermutation (Figure 1E), consistent with a life history lacking a stage of classical affinity maturation. If persistent lineages are indeed derived from B-1 cells, our results strongly suggest that expansion and maintenance of B-1 cell populations are homeostatic neutral processes with balanced birth and death rates, in sharp contrast with the strong positive selection that shapes vaccine-responsive B cells. B-1 cells exhibit preferential usage of V gene families (*27*), so selection on allelic diversity at the level of human populations may drive the evolution of these genes. The molecular identity of human B-1 cells has been elusive (*28*), and our prediction that these cells are distinguished by the genetic signatures of somatic evolution opens a new avenue for identification and characterization of this cell population.

How is the clonal structure of individual B cell lineages influenced by selection? During affinity maturation, subclones harboring independent mutations within a B cell lineage compete with one another for evolutionary success. Competition can result in either one winner or multiple winners within a clonal lineage. Multiple winners may arise due to independent competition in spatially separated regions, such as different GCs, or because subclones harboring different beneficial mutations compete to a stalemate within the same GC, a scenario known as “clonal interference” (*29*). To further dissect the evolutionary processes of affinity maturation, we characterized the clonal structures of vaccine-responsive lineages.

Using phylogenetic analysis, we found that many vaccine-responsive clonal B cell lineages contain multiple positively selected subclones. While some phylogenies harbor only one imbalanced clade displaying characteristics of recurrent selective sweeps (Figure 2A), others have several large clades that each exhibit these characteristics, suggesting that multiple subclones persisted as winners within these clonal lineages (Figure 3A). To quantify this phenomenon, we developed an algorithm to identify and count positively selected subclones in an unbiased manner (Materials and Methods). We found that 24% of vaccine-responsive lineages composed of >1000 sequences harbor multiple subclones that have evidence of positive selection (Figure 3B; false discovery rate of 1%). This indicates that affinity maturation often focuses the repertoire onto multiple subclones arising from a common B cell ancestor. These subclones share somatic mutations that were acquired prior to branching in every case, which is evidence against these results being artifacts arising from erroneous joining of non-clonal sequences during lineage reconstruction. The number of selective sweeps within a lineage is modestly but significantly correlated with the population size of the lineage (Figure 3C), suggesting that massive clonal amplification of vaccine-responsive B cell lineages often involves selection favoring multiple subclones. Previous reports indicate that clonally related sequences are occasionally found in distinct GCs located within the same lymph node (*13*), supporting a role for spatial segregation in facilitating independent selection of subclones.

**Figure 3.**
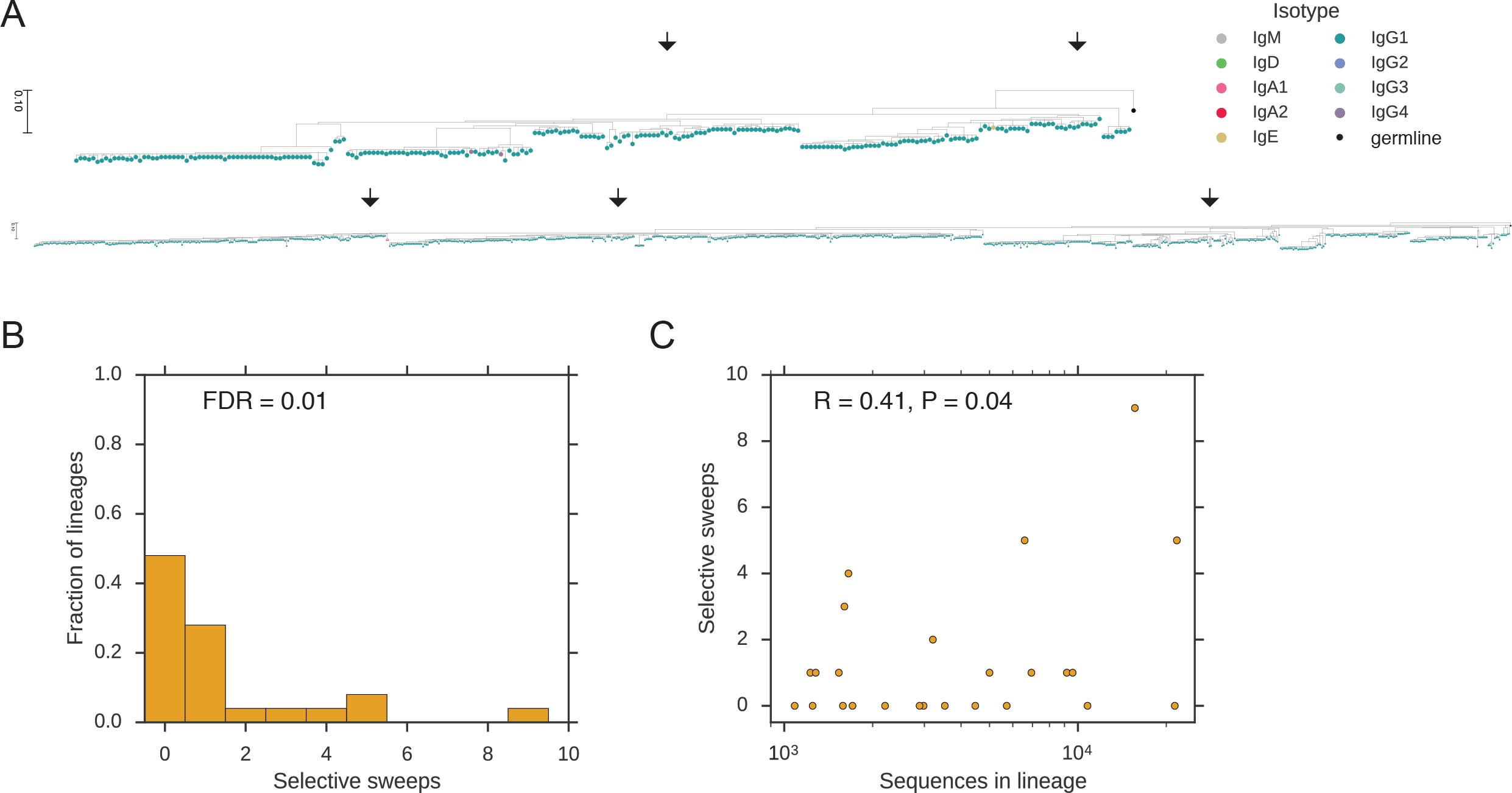
Signatures of selective sweeps within multiple subclones of vaccine-responsive antibody lineages. (A) Examples of phylogenies of vaccine-responsive clonal B cell lineages having evidence for selective sweeps favoring multiple subclones. Arrows indicate clades that were identified as significantly positively selected by our algorithm (P < 0.05). Leaves are colored by isotype. Phylogenies are rooted on the germline sequence. (B) Distribution of the number of distinct selective sweeps within vaccine-responsive lineages having >1,000 sequences. FDR, false discovery rate. (C) Relationship between the number of distinct selective sweeps within a clonal lineage and the population size (number of sequences) of the lineage. Pearson correlation coefficient is shown.

Because B cell fitness is tightly linked to antibody affinity during affinity maturation, we hypothesized that the genetic diversity of B cell populations encodes information about binding affinity. Amplification of highly fit variants can be readily observed in phylogenies, and elevated fitness is thought to be associated with enhanced antibody affinity. To test this idea, we sought to leverage phylogenetic signals that reveal the fitness of individual antibody sequences in order to rapidly identify candidate high-affinity antibodies and affinity-enhancing mutations based on sequencing data alone. Specifically, we used a computational approach to infer the fitness of sequences based on their phylogenetic context (*30*) and then identified specific sequences that had high fitness.

In line with a history of selective sweeps, phylogenetic inference revealed wide variation in fitness among sequences within vaccine-responsive B cell lineages, with some sequences predicted to have much higher fitness than other sequences in the same clonal lineage (Figure 4A). We identified mutations associated with the strongest fitness enhancements (top 3 branches ranked by fitness change from parent to child sequence in each lineage) (Materials and Methods). In comparison with synonymous mutations, non-synonymous fitness-enhancing mutations were highly enriched in CDRs (Figure 4B; P < 0.008 for CDR1, P < 0.1 for CDR2, and P < 2 × 10^−6^ for CDR3; Fisher’s exact test, twosided) and depleted in FWRs (P < 0.009 for FWR1, P < 2 × 10^−11^ for FWR3, and P < 0.01 for FWR4) with the sole exception of FWR2 (P = 0.87). This finding supports the functional relevance of the identified fitness enhancement-associated non-synonymous mutations and is consistent with the structural basis of antibody-antigen interactions (*31–33*). Mutations associated with the strongest fitness diminishments (bottom 3 branches in each lineage) were also enriched in CDR3 (Figure 4B; P < 8 × 10^−11^), which is consistent with the idea that mutations in CDRs can sometimes harm fitness because they disrupt antibody-antigen binding interfaces, suggesting that the traditional notion of purifying selection being confined to FWRs is overly simplistic. Together these results show how phylogenetic methods can potentially reveal information about antibody affinity that is encoded in sequence diversity and thus be used to identify high-affinity antibodies and affinity-enhancing mutations.

**Figure 4.**
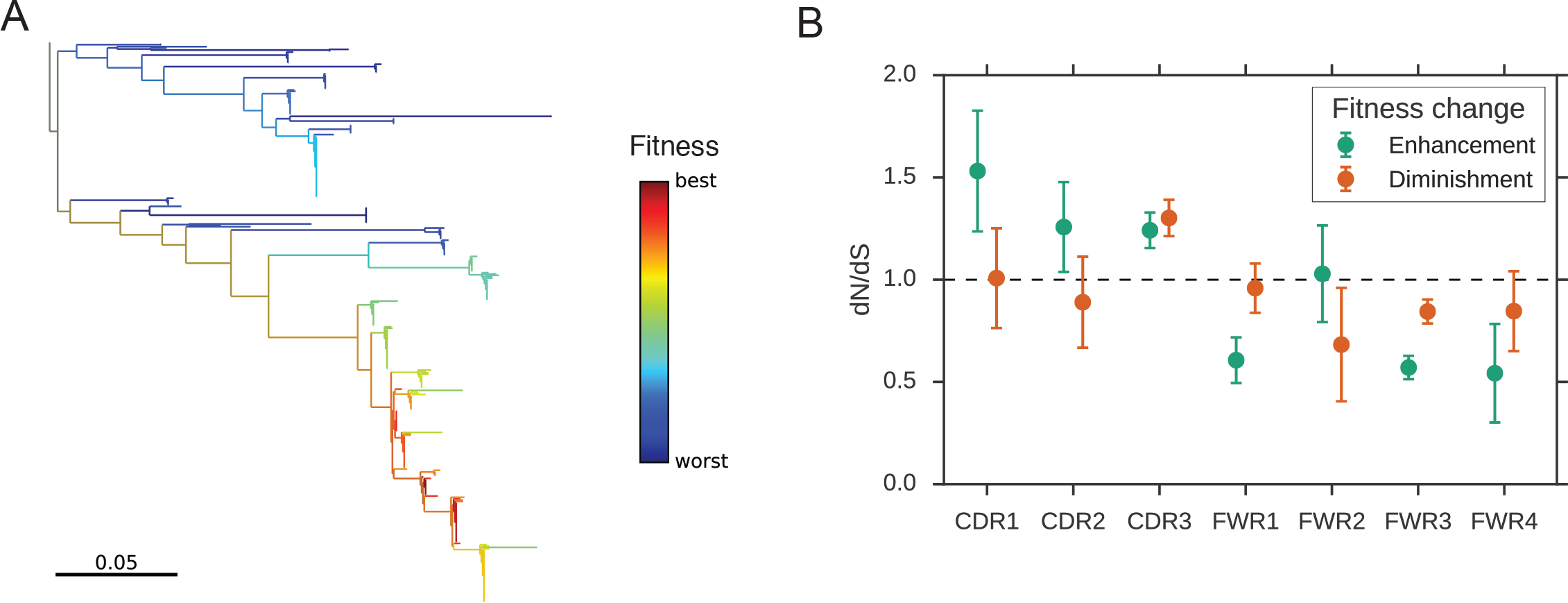
Phylogenetic identification of affinity-enhancing mutations. (A) Example of a phylogeny of a clonal B cell lineage colored by the inferred fitness of each sequence. (B) Regional distribution of non-synonymous mutations associated with strong fitness enhancements (top 3 branches ranked by fitness change from parent to child) or diminishments (bottom 3 branches ranked by fitness change from parent to child), displayed as enrichment relative to synonymous mutations (dN/dS) in the same branches. Dashed line indicates no enrichment. Error bars indicate one standard deviation as determined by bootstrap (100 replicates).

In summary, our results demonstrate that human antibody repertoires are shaped by a broad spectrum of somatic evolutionary processes. Prior efforts to detect selection in antibody genes have focused on regions or residues in aggregate across many clonal B cell lineages (*23*, *24*, *34*), and did not account for the fact that evolution acts differently on different clonal lineages. We have characterized signatures of selection within individual clonal B cell lineages up to the fundamental limits imposed by their population size, revealing that a diversity of evolutionary modes exists within the B cell repertoire. Vaccine-responsive lineages display pervasive evidence of positive selection, and many lineages experience selective sweeps favoring multiple subclones, revealing how the complex clonal structure of B cell populations is shaped by selection during affinity maturation. On the other hand, persistent lineages display signatures of neutral drift-like evolution, revealing that non-selective birth-death processes generate a substantial fraction of human antibody repertoires and requiring a major modification of the conventional notion that selective processes are ubiquitous in antibody maturation. This diversity of evolutionary modes likely reflects the diversity of life histories among distinct B cell types. The presence of large clonal lineages lacking molecular signatures of selection also provides an inherent control and constitutes evidence that the detection of such signatures in vaccine-responsive lineages is not an artifact of our approach, including a failure to correctly determine the germline sequence. Importantly, we have shown that the molecular signatures of selection distinguish the vaccine-responsive lineages from the persistent clonal lineages that are also highly abundant after vaccination. These signatures can also be harnessed through phylogenetic approaches to identify sequences that were most favored by selection during affinity maturation and thus are likely to encode high-affinity antibodies, demonstrating potential utility for biomedical applications. High-throughput sequencing of human antibody repertoires and analysis through the lens of population genetics therefore offers a promising avenue for antibody discovery and engineering.

## Acknowledgments

We thank our study volunteers for their participation. We thank Sally Mackey for Regulatory and Clinical Project Management, Research Nurses Susan Swope and Tony Trela, and Phlebotomist Michele Ugur for conducting the study visits. We thank Lily Blair, Elizabeth Jerison, Fabio Zanini, Derek Croote, David Glass, Richard Neher, and Jonathan Pritchard for discussions. This work was supported by NIH U19A1057229 (S.R.Q.) and the National Science Foundation Graduate Research Fellowship Program (F.H.). The clinical project was supported by NIH/NCRR CTSA award number UL1 RR025744 and clinical trial information is available from ClinicalTrials.gov (identifier NCT02987374). Sequence data is available from the Sequence Read Archive (accession number XXX). Preprocessed data are available at XXX and code is available at https://github.com/felixhorns/BCellSelection. The authors declare no competing interests.

## Supplementary Materials

**Figure S1.**
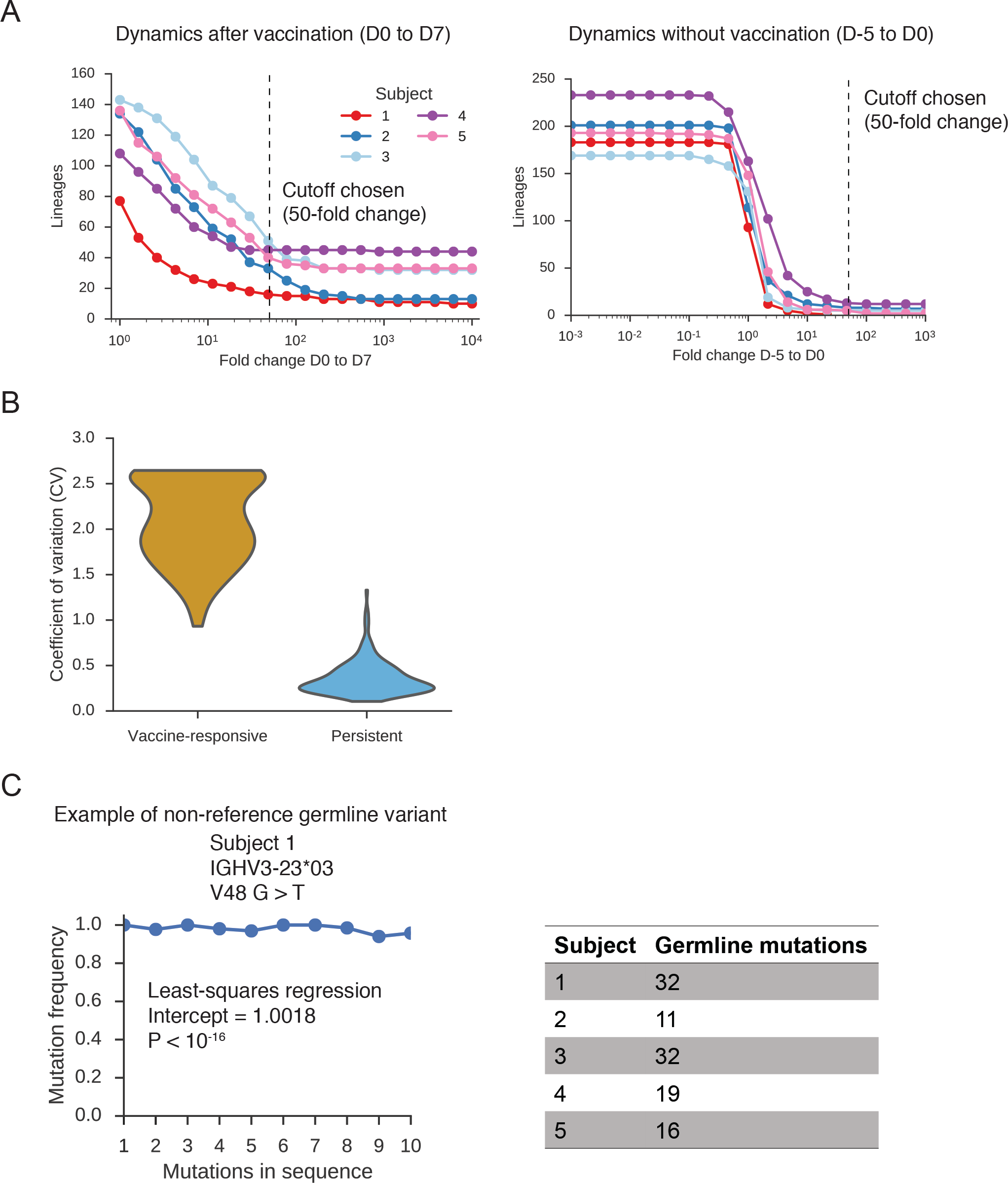
Dynamics of antibody repertoires and personalized annotation of germline variants. (A) Effect of cutoff for identifying vaccine-responsive lineages. Plots show the number of lineages having a significant change in abundance as a function of the fold-change cutoff used to determine significance. Right panel, comparison of D-5 to D0 (no vaccination). Left panel, comparison of D0 to D7 (after vaccination). We chose the cutoff of >50-fold change because at this value few lineages (27 among all five subjects) are identified as having a significant change in abundance in the absence of vaccination (D-5 to D0). The abundance fluctuation of these lineages may be due to environmental exposure to antigens, and in fact most of these lineages had infinite fold-change on the interval D-5 to D0 because they were not detected at D-5. The identity of vaccine-responsive lineages is largely insensitive to the choice of fold-change cutoff across a broad range (10-fold to 10,000-fold increase) because most vaccine-responsive lineages are not observed at D0 and therefore have infinite fold-change on the interval D0 to D7. (B) Dynamical variation in the fractional abundance of vaccine-responsive and persistent lineages. Plot shows the distribution of the coefficient of variation of fractional abundance for individual lineages across the observation period. (C) Personalized annotation of germline variants for study subjects using the method of Gadala-Maria and colleagues (*35*). Left panel shows an example of an identified nonreference germline variant. Identification is based on the presence of a y-intercept value that is significantly larger than zero. Right panel shows the number of non-reference germline variants detected for each subject.

**Figure S2.**
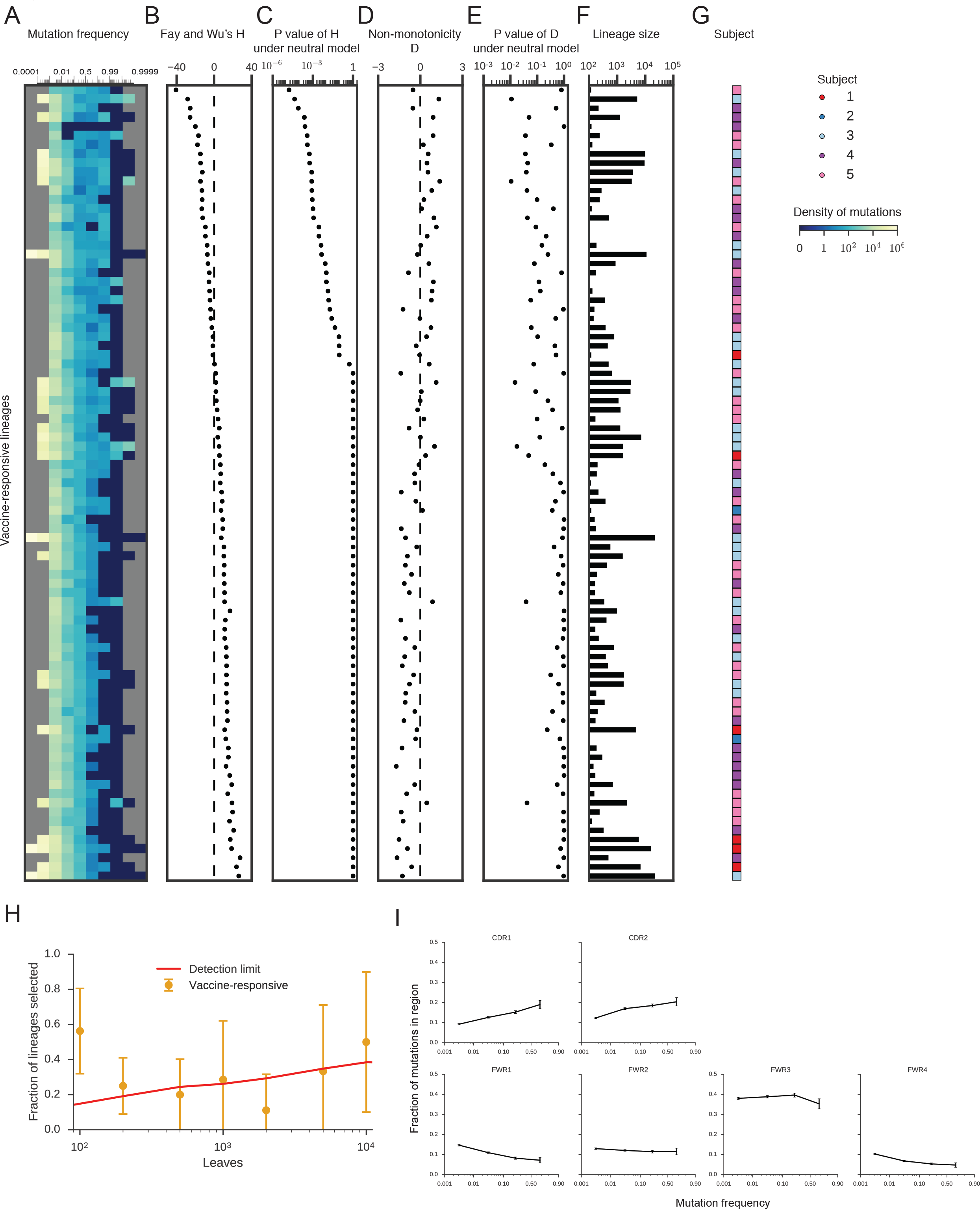
Genetic signatures of selection in individual vaccine-responsive B cell lineages. (A) Site frequency spectrums (SFSs) of individual clonal vaccine-responsive B cell lineages. The density of mutations in each frequency bin is indicated by color. (B) Fay and Wu’s H statistic of each lineage. (C) Significance of Fay and Wu’s H statistic in comparison with a null model of neutral drift. (D) Non-monotonicity D of the SFS of each lineage. (E) Significance of the non-monotonicity D in comparison with a null model of neutral drift. In (C) and (E), significance values were calculated by creating a ensemble of lineages via simulation of the Kingman coalescent model (neutral drift-like evolution) with each lineage having a population size matching that of the focal lineage, calculating the desired test statistic on each simulated lineage, fitting the Johnson’s U distribution to the simulated distribution of test statistics, then calculating the P value of the observed value of the test statistic. (F) Number of sequences in each lineage observed at D7 using long amplicon sequencing (paired-end 300 bp sequencing of 480 bp amplicons). (G) Subject of origin of each clonal B cell lineage. (H) Rate of detecting selection among clonal lineages of varying size. Detection limit imposed by population size is shown for comparison, assuming a false discovery rate of 0.05. Error bars show exact binomial 95% confidence intervals. (I) Distribution of mutations across sequence regions for mutations of different frequencies found in vaccine-responsive lineages. All mutations were placed into bins based on their frequency, then within each bin the fraction of mutations falling in each region was calculated. Error bars show exact binomial 95% confidence intervals.

**Figure S3.**
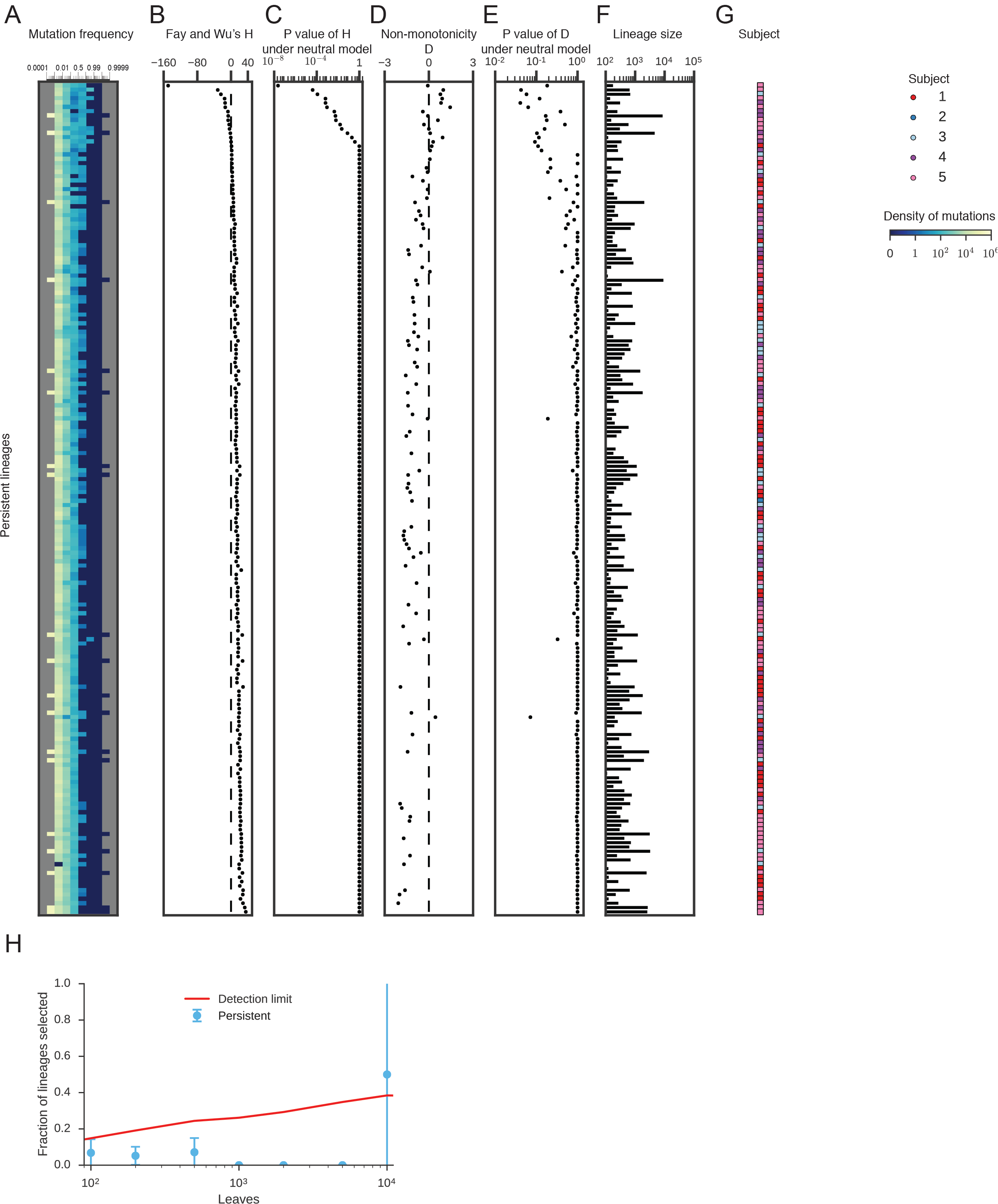
Genetic signatures of neutral evolution in individual persistent B cell lineages. (A) Site frequency spectrums (SFSs) of individual clonal persistent B cell lineages. The density of mutations in each frequency bin is indicated by color. (B) Fay and Wu’s H statistic of each lineage. (C) Significance of Fay and Wu’s H statistic in comparison with a null model of neutral drift. (D) Non-monotonicity D of the SFS of each lineage. (E) Significance of the non-monotonicity D in comparison with a null model of neutral drift In (C) and (E), significance values were calculated as described in Figure S2. (F) Number of sequences in each lineage observed at D7 using long amplicon sequencing (paired-end 300 bp sequencing of 480 bp amplicons). (G) Subject of origin of each clonal B cell lineage. (H) Rate of detecting selection among clonal lineages of varying size. Detection limit imposed by population size is shown for comparison, assuming a false discovery rate of 0.05. Error bars show exact binomial 95% confidence intervals.

**Figure S4.**
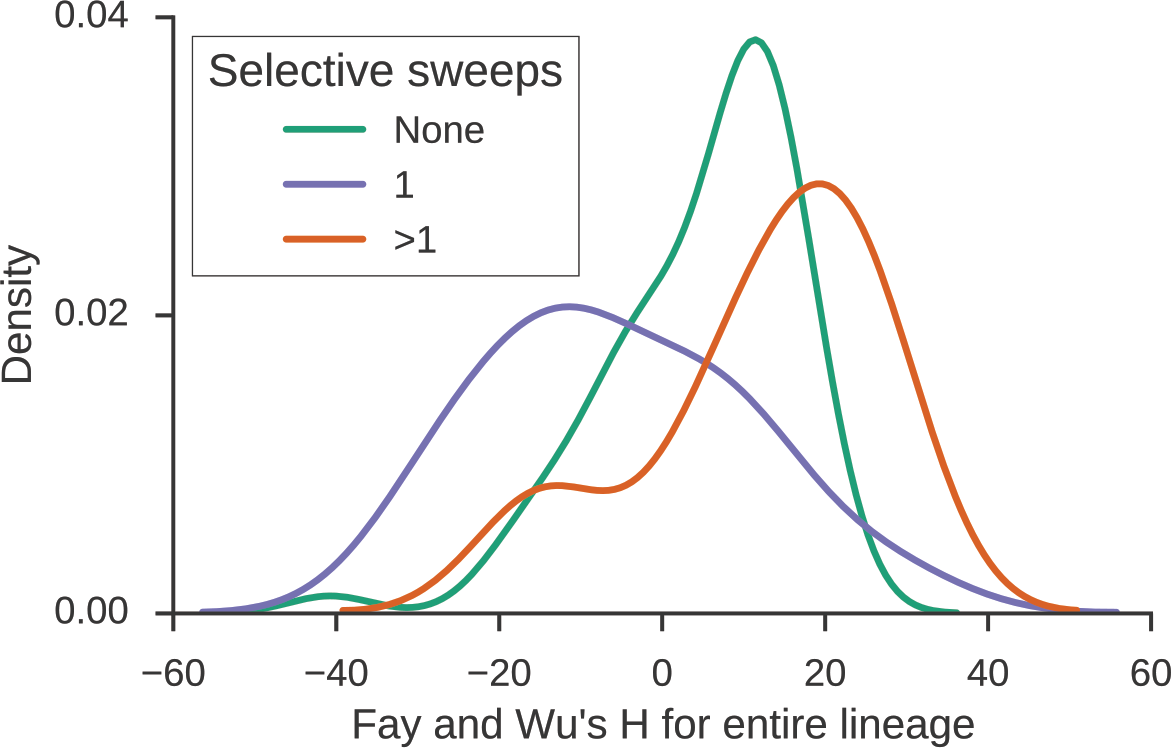
Effect of selective sweeps favoring multiple subclones on detection of selection. Distributions of Fay and Wu’s H statistic for vaccine-responsive lineages in which one, multiple, or no subclones have evidence for positive selection (false discovery rate of 1%). Lineages where multiple subclones were selected display a distinct rightward shift in the distribution of H, reflecting the hard upper bound on the frequency of mutations that are private to each subclone and causing the failure of this test for selection when applied to the entire lineage.

**Table S1.** Vaccine-responsive and persistent lineages found in each subject.

| Subject | Vaccine-responsive lineages | Persistent lineages |
| --- | --- | --- |
| 1 | 16 | 111 |
| 2 | 32 | 97 |
| 3 | 49 | 44 |
| 4 | 45 | 76 |
| 5 | 40 | 89 |

**Table S2.**
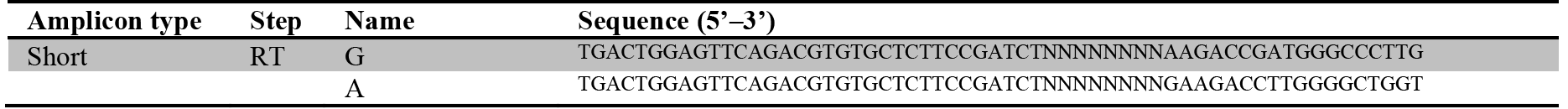
PCR primers used for library preparation.

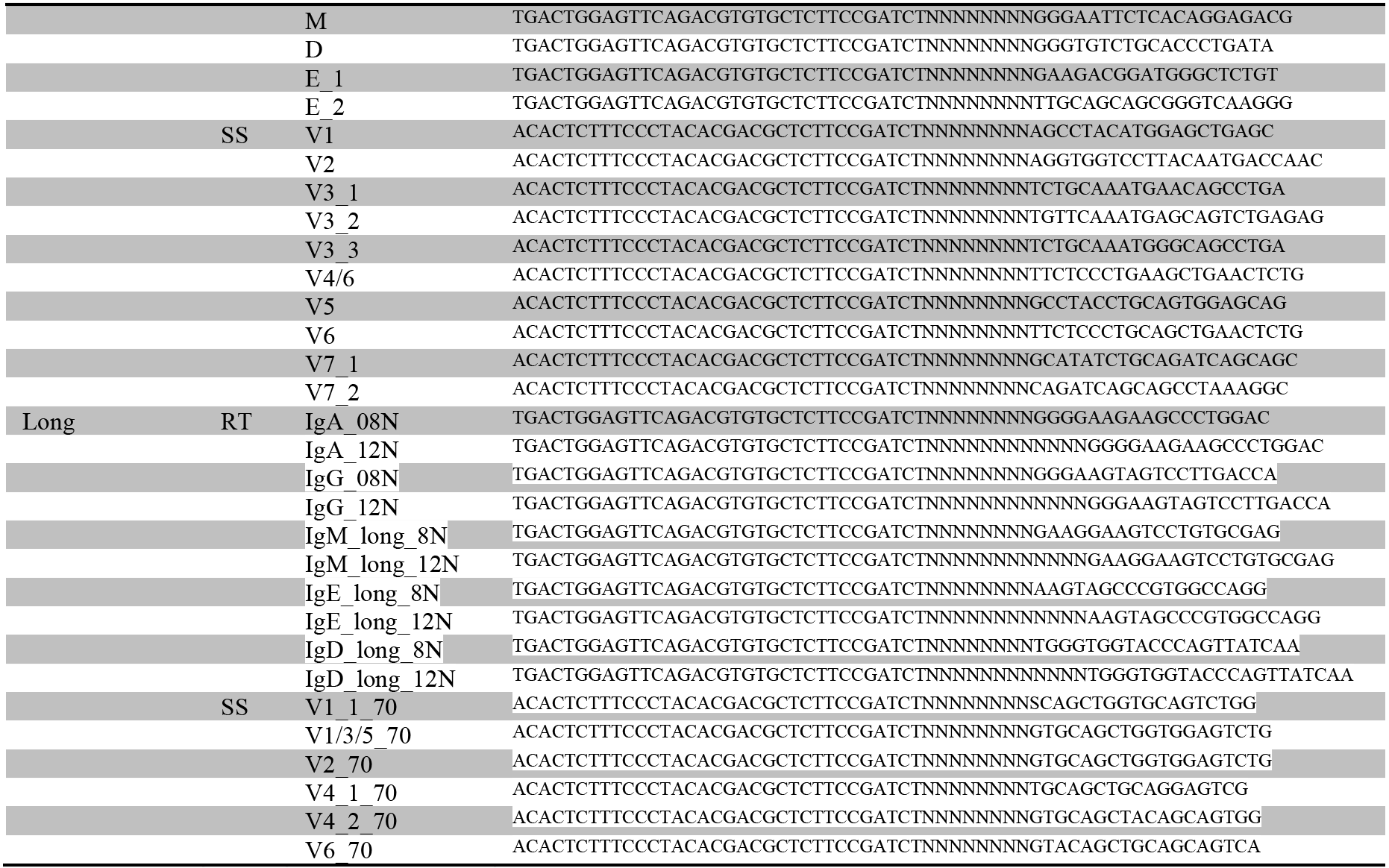

Short amplicon primers were used to prepare libraries for paired-end 100 bp sequencing from samples from all time points. Long amplicon primers were used to prepare libraries for paired-end 300 bp sequencing from samples from D7. RT, reverse transcription; SS, second-strand synthesis.

## Materials and Methods

### Study participants

All study participants gave informed consent and protocols were approved by the Stanford Institutional Review Board. Five humans aged 18-28, including 3 males and 2 females, were recruited in 2011. All subjects were apparently healthy and showed no signs of disease.

### Sample collection

Blood was drawn by venipuncture. Peripheral blood mononuclear cells (PBMCs) were isolated using a Ficoll gradient and frozen in 10% (vol/vol) DMSO and 40% fetal bovine serum (FBS) according to Stanford Human Immune Monitoring Center protocol. Subjects were vaccinated with the 2011–2012 seasonal trivalent inactivated influenza vaccine. Blood was collected 3 and 5 days before vaccination (D-3 and D-5); immediately before vaccination (D0); and 1, 4, 7, 9, and 11 days afterwards (D1, D4, D7, D9, D11).

### RNA extraction and library_preparation

PBMCs were thawed on ice and total RNA was extracted using the Qiagen AllPrep kit (Valencia, CA) following manufacturer’s instructions. Sequencing libraries were prepared from samples at all time points using 500 ng of total RNA as input following the protocol described in (*14*). Briefly, RNA was annealed to a pool of six isotype-specific IGH constant region primers containing 8 random nucleotides (nt), which serve as a molecular barcode for consensus error correction, by incubation at 72 C for 3 min, then placed on ice for 2 min. First-strand cDNA synthesis was performed using Superscript III reverse transcriptase (Life Technologies, Carlsbad, CA) following manufacturer’s protocol. Second-strand cDNA synthesis was performed using Phusion HiFi DNA polymerase (Thermo Scientific, Waltham, MA) and a pool of ten IGH variable region-specific primers containing 8 random nt (98°C for 4 min, 52°C for 1 min, 72°C for 5 min). Double-stranded cDNA product was purified twice using Ampure XP beads (1:1 ratio) (Beckman Coulter, Indianapolis, IN). Amplification was performed using Platinum Hifi DNA polymerase (Life Technologies, Carlsbad, CA) and primers containing Illumina sequencing adapters and dual sample indexes. Products were purified using Ampure XP beads (1:1 ratio), then pooled for multiplexed sequencing.

We prepared additional sequencing libraries from D7 following the protocol described in (*15*). This protocol is identical to that described above, except that ten isotype-specific IGH constant region primers and six IGH variable region-specific primers each containing 8 or 12 random nt were used. Products are longer amplicons spanning most of the IGH variable region and ~100 bp of the IGH constant region. We used a different aliquot of total RNA from the same D7 samples as input. All PCR primer sequences are provided in Table S2.

### Sequencing

Sequencing was performed for libraries from all time points using the Illumina HiSeq 2500 platform (San Diego, CA) using paired-end 101 bp reads. For libraries prepared from the D7 time point with longer amplicons, sequencing was performed using the Illumina Miseq platform using paired-end 300 bp reads. We obtained 826472 ± 413841 reads (mean ± s.d., range 170477 – 1988165) for each library.

### Preprocessing of sequence data

To process sequencing reads, we used a custom informatics pipeline similar to (*15*). Briefly, consensus sequences were constructed from reads containing the same 16 nt random barcode. Quality scores were propagated to the consensus sequence. Sequences were annotated with V and J germline gene usage and CDR3 length using IgBlast (36). Isotypes were determined using BLASTN against a custom database of IGH constant region sequences.

### Identification of clonal B cell lineages

Sequences belonging to the same clonal B cell lineage were identified using clustering following (*15*). Briefly, sequences sharing the same V and J germline genes and CDR3 length were grouped. Within each group, single-linkage clustering was performed with a cutoff of 90% nt sequence identity across both the CDR3 and the rest of the variable region. Sequence identity was computed by counting mismatches in gapless pairwise sequence alignments. Quality filtering was implemented by assuming mismatches at positions where either aligned base had Q ≤ 5. The cutoff of 90% was chosen because it is a distinct minimum in the distribution of pairwise nucleotide distances between sequences (data not shown). This approach has been shown to partition sequences into clonal lineages with high sensitivity and specificity (*15*, *16*).

### Tracking dynamics of clonal B cell lineages

To track the dynamics of clonal B cell lineages, we calculated the fractional abundance of the each lineage, defined as the number of unique sequences within the lineage divided by the total number of unique sequences observed in the repertoire at that time point. For this calculation, we only used reads that were sequenced using the short amplicon protocol. Vaccine-responsive lineages were identified based on the fold-change of their fractional abundance between D0 and D7 (>50-fold increase). Persistent lineages were identified as those having stable fractional abundance between D0 and D7 (<2-fold increase). We further required that each vaccine-responsive and persistent lineage represent >0.1% of the repertoire at D7 (corresponding to ~40 distinct sequences) to remove very small clonal lineages from consideration. Isotype composition and mutation density were calculated using sequences from all time points.

### Identification of non-reference germline variants

To annotate non-reference germline variants in a personalized manner for each subject, we adapted the method proposed by Gadala-Maria and colleagues (*35*). We first grouped sequences having the same V or J germline sequence. For each mutation detected by comparison against the reference germline sequence, we performed regression on the mutation frequency against the mutation count of the entire V or J segment. Specifically, we binned sequences into groups based on the number of mutations per sequence and calculated the frequency of the focal mutation in each bin. We then fit a linear model to these data using least-squares optimization. Mutations with y-intercepts greater than 0.125 at a significance level of P < 0.05 as assessed using Student’s t test were considered potential germline variants. Because alleles might contain multiple nonreference germline variants, bins were excluded from the regression based on detection of outliers (bins having more than 1.5-times the interquartile range greater than the third quartile of the number of sequences in the bins carrying 1–10 mutations). If an outlier bin was found, then all bins having fewer mutations per sequence were excluded from the regression.

### Calculation of site frequency spectrums

We constructed the site frequency spectrum (SFS) of each clonal B cell lineage based on somatic mutations relative to the germline V and J genes. For analysis of the SFS and further phylogenetic analysis, we used only reads originating from the D7 samples that were sequenced using the long amplicon protocol. Vaccine-responsive and persistent lineages having <100 unique sequences in these samples were excluded from this analysis. Mutations were called using IgBlast (*36*) and we removed non-reference germline variants for each individual subject as determined above. We calculated the frequency of each mutation within a clonal lineage (number of sequences containing the mutation / number of sequences in the lineage). We note that our approach conservatively excludes most mutations in the CDR3 because these mutations lie within the highly variable untemplated region of the IGH sequence and therefore the ancestral state may not be known with high confidence.

To visualize the SFS, we binned the mutation frequencies using bins spaced according to the logit function (inverse logistic transform). Bin edges were 10^−5^, 10^−4^, 10^−3^, 10^−2^, 10^−1^, 0.5, 0.9, 0.99, 0.999, 0.9999, 0.99999. The mutation density within each bin was calculated by normalizing by the bin size (number of mutations in bin / width of bin). To calculate the average SFS across many lineages (e.g., all vaccine-responsive lineages or vaccine-responsive lineages from one study subject), we calculated the SFS for each lineage individually, then calculated the average mutation density in each bin. Each lineage is weighted equally and therefore the average is not influenced by the population sizes or relative mutational loads of the lineages.

Use of the SFS for detecting selection has several practical advantages. Calculation of the SFS does not depend on phylogenetic reconstruction or ancestral sequence reconstruction and the reliability of these inferences. Unlike traditional tree imbalance measures, such as the Colless or Sackin indices, the SFS is readily calculated for populations with multifurcating phylogenies, such as B cell populations. Finally, calculation of the SFS scales linearly with the number of sequences and therefore can be evaluated readily for lineages having many sequences.

### Simulations of evolutionary models

To compare the observed patterns of evolution with evolutionary models, we performed simulations of beta coalescent models using the betatree package in Python (*37*). Specifically, we simulated neutral evolution using the Kingman coalescent (α = 2) and evolution under strong positive selection using the Bolthausen-Sznitman coalescent (α = 1). For comparison with the observed SFSs averaged across many lineages, we simulated ensembles of 100 lineages (similar to the number of observed vaccine-responsive lineages) each having a number of leaves sampled without replacement from the distribution of population sizes of vaccine-responsive lineages (median population size was approximately 1,000 sequences), and calculated the average SFS across these lineages (Figure 2A and Figure 2B).

### Calculation of test statistics for selection

We calculated Fay and Wu’s H statistic on the counts of somatic mutations within a clonal lineage:

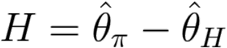

with

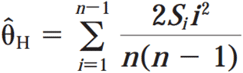
and

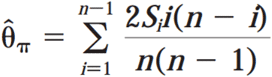
where S_i_ is the number of mutations observed in *i* sequences of the lineage and *n* is the total number of sequences in the lineage, i.e. the population size of the lineage (*22*).

As an alternative metric for selection, we directly estimated the non-monotonicity of the high-frequency region of the SFS. Specifically, we fit a quadratic polynomial to the binned SFS using least-squares minimization, calculated its first derivative, and determined the maximum value of the first derivative in bins representing frequencies >0.25, which we define as the non-monotonicity *D*. SFSs having an excess of high-frequency mutations display a characteristic “uptick” or non-monotonicity in the high-frequency region and therefore have positive *D*.

### Calculation of the statistical significance of test statistics

We evaluated the statistical significance of tests for selection by comparison with a null distribution of the test statistic generated under a neutral model of evolution (the Kingman coalescent). We simulated an ensemble of 1,000 lineages using the Kingman coalescent and calculated the test statistic (Fay and Wu’s H or the non-monotonicity *D*) for each lineage. Thus, we created a distribution of the test statistic under the null model. We then fitted the Johnson’s U distribution to this data. To evaluate the statistical significance of a test statistic for a focal lineage, we calculated the P value of the test statistic (that is, the probability of obtaining by chance a value of the test statistic that is at least as extreme as the given value) under the null distribution by integrating its probability density. Because population size strongly influences the distribution of test statistics, we always tested for selection by comparison against a null distribution characterizing populations of a size matched to that of the focal lineage. To accomplish this, we simulated the null distribution as described above for a range of population sizes (N = 100, 200, 500, 1000, 2000, 5000, 10000, and 20000 leaves). Given a focal lineage, we determined the nearest lineage size within this set and used the corresponding null distribution for comparison. We refer to this procedure as matching the population size of the focal lineage to the null distribution.

### Determining the limit of detection of selection due to population size

Detection of selection is fundamentally limited by population size. The detection limit was calculated by simulating an ensemble of 1,000 lineages under strong positive selection using the Bolthausen-Sznitman coalescent model. Fay and Wu’s H statistic was calculated for each lineage and its significance was assessed by comparison with the neutral model. This was repeated for populations having various sizes (N = 100, 200, 500, 1000, 2000, 5000, 10000, 20000 leaves). The fraction of lineages that were identified as significantly positively selected (P < 0.05) in each case is the expected rate of detecting positive selection in the scenario where all lineages are generated under strong positive selection.

### Phylogenetic reconstruction

We used a fast heuristic algorithm to construct a multiple sequence alignment and reconstruct the phylogeny of each clonal lineage. Sequences were first aligned in an ungapped manner using the start and end positions of the CDR3 as anchor points. This alignment was refined using MUSCLE with “-refine – maxiters 1 – diags – gapopen -5000” (38). The large gap penalty reflects our expectation that insertions and deletions are uncommon during somatic hypermutation (*24*, *39*). We aligned a germline sequence consisting of the concatenated V and J germline alleles of the lineage by profile-profile alignment using MUSCLE with “-profile – maxiters 1 – diags”. We reconstructed the phylogeny using FastTree 2 with “-nt – gtr” (*40*). Finally, we performed joint refinement of the multiple sequence alignment and phylogeny by identifying extremely long branches (>0.5 substitutions/site), removing them from the alignment, and realigning one sequence at a time by profile-profile alignment using MUSCLE with “-profile – maxiters 2 – diags”, then repeating phylogenetic reconstruction as described above.

### Detecting selection in multiple subclones of a clonal lineage

We developed an algorithm to identify subclones having evidence of positive selection. Our algorithm is based on calculation of the test statistic on subclones, then searching within the phylogeny to identify the largest independent subclones displaying significant evidence of selection. Specifically, we calculate Fay and Wu’s H statistic on every large clade (having >100 sequences) based on the frequency of somatic mutations that occurred within the clade, and calculate its P value by comparison with the null distribution for phylogenies matched in size to the number of leaves in the clade. We then perform a greedy breadth-first search for clades having significant evidence of selection. This search strategy yields the deepest subclones having evidence of selection and guarantees that all such subclones represent mutually exclusive subsets of the lineage. We note that this is a conservative strategy because in a case where a deep clade has evidence of selection, but in turn harbors two independent subclades that themselves have evidence of selection, the search stops at the deep clade and therefore will not discover the selected subclades. To correct for multiple hypothesis testing, we adjusted the P value associated with Fay and Wu’s H statistic using the Bonferroni method based on the number of tests performed during the search step.

We observed that standard tests of selection, such as Fay and Wu’s H statistic, often failed to detect selection when applied to lineages harboring multiple positively selected subclones. When multiple subclones persist, the frequency of a derived mutation which is private to a single subclone has a hard upper bound, causing tests based on the presence of high frequency mutations to fail (Figure S4). This highlights the influence of clonal population structure on tests for selection, an important design consideration for efforts to detect selection in any asexual population.

### Identification of candidate affinity-increasing mutations

Using the reconstructed phylogeny of each clonal lineage as input, we performed fitness inference following (*30*). Fitness inference is based on the idea that nodes having higher fitness create offspring at a faster rate than other nodes and therefore the local branching rate of a phylogeny carries information about the fitness of sequences within the phylogeny. Fitness was inferred using fitness diffusion constant D = 0.5, distance scale = 2.0, and sampling fraction = 0.1. We annotated each branch with the mean fitness change from the parent to the child node. To identify branches having large fitness enhancements or diminishments, we ranked all branches by their fitness change and selected those among the top 3 or bottom 3. Our conclusions also hold true when analysis is performed using the top and bottom 1, 5, or 10 branches. We performed ancestral sequence reconstruction for each clonal lineage using maximum-likelihood assuming equal rates for all mutations. We then identified mutations that occurred on each branch by comparing the reconstructed parent and child sequences. We assigned these mutations to regions (CDRs and FWRs) based on the region boundaries identified using IgBlast (*36*). To compute the enrichment of non-synonymous mutations in a region in comparison with synonymous mtuations (dN/dS), we calculated the fraction of non-synonymous mutations falling in a region, and then divided this fraction by the corresponding fraction calculated using synonymous mutations. We calculated the error of this measurement by bootstrap resampling of branches (100 replicates).

## References and Notes

1. F. M. Burnet, A Modification of Jerne’s Theory of Antibody Production using the Concept of Clonal Selection. Australian J. Sci. 20, 67–9 (1957).

2. H. N. Eisen, Affinity Enhancement of Antibodies: How Low-Affinity Antibodies Produced Early in Immune Responses Are Followed by High-Affinity Antibodies Later and in Memory B-Cell Responses. Cancer Immunol Res. 2, 381–392 (2014).

3. D. M. Tarlinton, Evolution in miniature: selection, survival and distribution of antigen reactive cells in the germinal centre. Immunol. Cell Biol. 86, 133–138 (2008).

4. G. D. Victora, P. C. Wilson, Germinal Center Selection and the Antibody Response to Influenza. Cell. 163, 545–548 (2015).

5. H. N. Eisen, G. W. Siskind, Variations in affinities of antibodies during the immune response. Biochemistry. 3, 996–1008 (1964).

6. M. Kuraoka et al., Complex Antigens Drive Permissive Clonal Selection in Germinal Centers. Immunity. 44, 542–552 (2016).

7. C. D. C. Allen, T. Okada, J. G. Cyster, Germinal-center organization and cellular dynamics. Immunity. 27, 190–202 (2007).

8. G. D. Victora, L. Mesin, Clonal and cellular dynamics in germinal centers. Current Opinion in Immunology. 28, 90–96 (2014).

9. G. D. Victora et al., Germinal Center Dynamics Revealed by Multiphoton Microscopy Using a Photoactivatable Fluorescent Reporter. Cell. 143, 592–605 (2010).

10. A. D. Gitlin, Z. Shulman, M. C. Nussenzweig, Clonal selection in the germinal centre by regulated proliferation and hypermutation. Nature. 509, 637–640 (2014).

11. A. D. Gitlin et al., T cell help controls the speed of the cell cycle in germinal center B cells. Science. 349, 643–646 (2015).

12. Z. Shulman et al., T Follicular Helper Cell Dynamics in Germinal Centers. Science. 341, 673–677 (2013).

13. J. M. J. Tas et al., Visualizing antibody affinity maturation in germinal centers. Science, aad3439 (2016).

14. C. Vollmers, R. V. Sit, J. A. Weinstein, C. L. Dekker, S. R. Quake, Genetic measurement of memory B-cell recall using antibody repertoire sequencing. PNAS. 110, 13463–13468 (2013).

15. F. Horns et al., Lineage tracing of human B cells reveals the in vivo landscape of human antibody class switching. eLife Sciences. 5, e16578 (2016).

16. N. T. Gupta et al., Hierarchical Clustering Can Identify B Cell Clones with High Confidence in Ig Repertoire Sequencing Data. J. Immunol. 198, 2489–2499 (2017).

17. T. Bedford, S. Cobey, M. Pascual, Strength and tempo of selection revealed in viral gene genealogies. BMC Evolutionary Biology. 11, 220 (2011).

18. F. Zanini et al., Population genomics of intrapatient HIV-1 evolution. eLife Sciences. 4, e11282 (2015).

19. R. A. Neher, O. Hallatschek, Genealogies of rapidly adapting populations. Proceedings of the National Academy of Sciences. 110, 437–442 (2013).

20. J. F. C Kingman, The coalescent. Stochastic Processes and their Applications. 13, 235–248 (1982).

21. E. Bolthausen, A.-S. Sznitman, On Ruelle’s Probability Cascades and an Abstract Cavity Method. Communications in Mathematical Physics. 197, 247–276 (1998).

22. J. C. Fay, C.-I. Wu, Hitchhiking Under Positive Darwinian Selection. Genetics. 155, 1405–1413 (2000).

23. G. Yaari, M. Uduman, S. H. Kleinstein, Quantifying selection in high-throughput Immunoglobulin sequencing data sets. Nucleic Acids Res. 40, e134–e134 (2012).

24. C. O. McCoy et al., Quantifying evolutionary constraints on B-cell affinity maturation. Phil. Trans. R. Soc. B. 370, 20140244 (2015).

25. I. Förster, K. Rajewsky, Expansion and functional activity of Ly-1+ B cells upon transfer of peritoneal cells into allotype-congenic, newborn mice. Eur. J. Immunol. 17, 521–528 (1987).

26. F. G. Kroese, W. A. Ammerlaan, A. B. Kantor, Evidence that intestinal IgA plasma cells in mu, kappa transgenic mice are derived from B-1 (Ly-1 B) cells. Int. Immunol. 5, 1317–1327 (1993).

27. T. D. Quách et al., Distinctions among Circulating Antibody-Secreting Cell Populations, Including B-1 Cells, in Human Adult Peripheral Blood. The Journal of Immunology. 196, 1060–1069 (2016).

28. N. Baumgarth, The double life of a B-1 cell: self-reactivity selects for protective effector functions. Nat Rev Immunol. 11, 34–46 (2011).

29. H. J. Muller, Some Genetic Aspects of Sex. The American Naturalist. 66, 118–138 (1932).

30. R. A. Neher, C. A. Russell, B. I. Shraiman, Predicting evolution from the shape of genealogical trees. eLife Sciences. 3, e03568 (2014).

31. E. A. Padlan et al., Structure of an antibody-antigen complex: crystal structure of the HyHEL-10 Fab-lysozyme complex. PNAS. 86, 5938–5942 (1989).

32. R. M. MacCallum, A. C. R. Martin, J. M. Thornton, Antibody-antigen Interactions: Contact Analysis and Binding Site Topography. Journal of Molecular Biology. 262, 732–745 (1996).

33. J. L. Xu, M. M. Davis, Diversity in the CDR3 Region of VH Is Sufficient for Most Antibody Specificities. Immunity. 13, 37–45 (2000).

34. I. S. Lossos, R. Tibshirani, B. Narasimhan, R. Levy, The inference of antigen selection on Ig genes. J. Immunol. 165, 5122–5126 (2000).

35. D. Gadala-Maria, G. Yaari, M. Uduman, S. H. Kleinstein, Automated analysis of high-throughput B-cell sequencing data reveals a high frequency of novel immunoglobulin V gene segment alleles. PNAS. 112, E862–E870 (2015).

36. J. Ye, N. Ma, T. L. Madden, J. M. Ostell, IgBLAST: an immunoglobulin variable domain sequence analysis tool. Nucleic Acids Res. 41, W34–W40 (2013).

37. R. A. Neher, T. A. Kessinger, B. I. Shraiman, Coalescence and genetic diversity in sexual populations under selection. PNAS. 110, 15836–15841 (2013).

38. R. C. Edgar, MUSCLE: multiple sequence alignment with high accuracy and high throughput. Nucl. Acids Res. 32, 1792–1797 (2004).

39. G. Teng, F. N. Papavasiliou, Immunoglobulin somatic hypermutation. Annu. Rev. Genet. 41, 107–120 (2007).

40. M. N. Price, P. S. Dehal, A. P. Arkin, FastTree 2 – Approximately Maximum-Likelihood Trees for Large Alignments. PLoS ONE. 5, e9490 (2010).

